# An AI Agent for Fully Automated Multi-omic Analyses

**DOI:** 10.1101/2023.09.08.556814

**Authors:** Juexiao Zhou, Bin Zhang, Xiuying Chen, Haoyang Li, Xiaopeng Xu, Siyuan Chen, Wenjia He, Chencheng Xu, Xin Gao

**Author notes:** These authors contributed equally to this work (Juexiao Zhou, Bin Zhang).

## Abstract

With the fast-growing and evolving omics data, the demand for streamlined and adaptable tools to handle the bioinformatics analysis continues to grow. In response to this need, we introduce Automated Bioinformatics Analysis (AutoBA), an autonomous AI agent designed explicitly for fully automated multi-omic analyses based on large language models. AutoBA simplifies the analytical process by requiring minimal user input while delivering detailed step-by-step plans for various bioinformatics tasks. Through rigorous validation by expert bioinformaticians, AutoBA’s robustness and adaptability are affirmed across a diverse range of omics analysis cases, including whole genome/exome sequencing (WGS/WES), chromatin immunoprecipitation assays with sequencing (ChIP-seq), RNA sequencing (RNA-seq), single-cell RNA-seq, spatial transcriptomics and so on. AutoBA’s unique capacity to self-design analysis processes based on input data variations further underscores its versatility. Compared with online bioinformatic services, AutoBA offers multiple LLM backends, with options for both online and local usage, prioritizing data security and user privacy. Moreover, different from the predefined pipeline, AutoBA has adaptability in sync with emerging bioinformatics tools. Overall, AutoBA represents an advanced and convenient tool, offering robustness and adaptability for conventional multi-omic analyses.

## 1 Introduction

Bioinformatics is an interdisciplinary field that encompasses computational, statistical, and biological approaches to analyze, understand and interpret complex biological data [1], [2], [3]. With the rapid growth of gigabyte-sized biological data generated from various high-throughput technologies, bioinformatics has become an essential tool for researchers to make sense of these massive datasets and extract meaningful biological insights. The applications of bioinformatics typically cover diverse fields such as genome analysis [4], [5], [6], structural bioinformatics [7], [8], [9], systems biology [10], data and text mining [11], [12], [13], phylogenetics [14], [15], [16], and population analysis [17], [18], which has further enabled significant advances in personalized medicine [19] and drug discovery [5].

In broad terms, bioinformatics could be categorized into two primary domains: the development of innovative algorithms to address various biological challenges [20], [21], [22], [23], [24], and the application of established tools to analyze extensive biological datasets [25], [26], especially high-throughput sequencing data. Developing new bioinformatics software requires a substantial grasp of biology and programming expertise. Alongside the development of novel computational methods, one of the most prevalent applications of bioinformatics is the investigation of biological data using the existing tools and pipelines [27], [28], which typically involves a sequential, flow-based analysis of omics data, encompassing variety types of datasets like whole genome sequencing (WGS) [29], whole exome sequencing (WES), RNA sequencing (RNA-seq) [30], single- cell RNA-seq (scRNA-Seq) [31], transposase-accessible chromatin with sequencing (ATAC-Seq) [32], ChIP-seq [33], and spatial transcriptomics [34].

For example, the conventional analytical framework for bulk RNA-seq involves a meticulously structured sequence of computational steps [35]. This intricate pipeline reveals its complexity through a series of carefully orchestrated stages. It begins with quality control [36], progresses to tasks such as adapter trimming [37] and the removal of low-quality reads, and then moves on to critical steps like genome or transcriptome alignment [38]. Furthermore, it extends to some advanced tasks, including the identification of splice junctions [39], quantification through read counting [40], and the rigorous examination of differential gene expression [41]. Moreover, the pipeline delves into the intricate domain of alternative splicing [42] and isoform analysis [43]. This progressive journey ultimately ends in downstream tasks like the exploration of functional enrichment [44], providing a comprehensive range of analytical pursuits. Compared to bulk RNA-seq, ChIP-seq involves distinct downstream tasks, such as peak calling [45], motif discovery [46], peak annotation [47] and so on. In summary, the analysis of different types of omics data requires professional skills and a comprehensive comprehension of the corresponding field, particularly for customized data analysis. Moreover, the methods and pipelines might vary across different bioinformaticians and they even may evolve with the development of more advanced algorithms.

Meanwhile, online bioinformatics analysis platforms are currently in vogue, such as iDEP [48], ICARUS [49] and STellaris [50]. However, they often necessitate the uploading of either raw data or pre-processed statistics by users, which could potentially give rise to additional privacy concerns and data leakage risks [51].

In the context described above, the bioinformatics community grapples with essential concerns regarding the standardization, portability, and reproducibility of analysis pipelines [52], [53], [54]. Moreover, achieving proficiency in utilizing these pipelines for data analysis demands additional training, posing challenges for many wet lab researchers due to its potential complexity and time-consuming nature. Even dry-lab researchers may find the repetitive process of running and debugging these pipelines to be quite tedious [55]. Meanwhile, bioinformatics data analysis training incurs substantial costs. The elevated expenses associated with training in bioinformatics data analysis could be attributed to the highly specialized nature of the field, the need for multi-modal data analysis, the evolution of technologies, restricted computing resources, the expense of training materials and tools, as well as the operational costs of training institutions. These factors collectively contribute to the high cost of bioinformatics training [56]. Consequently, there is a growing anticipation within the community for the development of a more user-friendly, low-code, multi-functional, automated, and natural language-driven intelligent tool tailored for end-to-end bioinformatics analysis. Such a tool has the potential to generate significant excitement and benefit researchers across the field.

**TABLE 1.** Summary of AutoBA application scenarios in bioinformatics multi-omics analysis. The table displays a comprehensive list of 40 real-world cases utilized to assess AutoBA, providing information on the class of the cases, the respective task name, and the corresponding case ID.

| Bioinformatics Pipelines | Tasks | Types of Omics | Case ID |
| --- | --- | --- | --- |
| WGS data analysis | Genome assembly | Genomics | 1.1 |
| WGS/WES data analysis | Somatic SNV+indel calling | Genomics | 2.1 |
| WGS/WES data analysis | Somatic SNV+indel calling and annotation | Genomics | 2.2 |
| WGS/WES data analysis | Structure variation identification with normal | Genomics | 2.3 |
| WGS/WES data analysis | Structure variation identification without normal | Genomics | 2.4 |
| ChIP-seq data analysis | Peak calling | Genomics | 3.1 |
| ChIP-seq data analysis | Motif discovery for binding sites | Genomics | 3.2 |
| ChIP-seq data analysis | Functional enrichment of target gene | Genomics | 3.3 |
| Bisulfite-Seq data analysis | Identifying DNA methylation | Genomics | 4.1 |
| ATAC-seq data analysis | Identifying open chromatin regions | Genomics | 5.1 |
| DNase-seq data analysis | Identifying DnaseI hypersensitive site | Genomics | 6.1 |
| 4C-seq data analysis | Find genomics interactions | Genomics | 7.1 |
| Nanopore DNA sequencing data analysis | Genome assembly | Genomics | 8.1 |
| Nanopore DNA sequencing data analysis | Tandem repeats variation identification | Genomics | 8.2 |
| PacBio DNA sequencing data analysis | Genome assembly | Genomics | 9.1 |
| RNA-Seq data analysis | Find Differentially expressed genes | Transcriptomics | 10.1 |
| RNA-Seq data analysis | Identify the top5 downregulated genes | Transcriptomics | 10.2 |
| RNA-Seq data analysis | Predict Fusion gene with annotation | Transcriptomics | 10.3 |
| RNA-Seq data analysis | Isoform expression | Transcriptomics | 10.4 |
| RNA-Seq data analysis | Splicing analysis | Transcriptomics | 10.5 |
| RNA-Seq data analysis | APA analysis | Transcriptomics | 10.6 |
| RNA-Seq data analysis | RNA editing | Transcriptomics | 10.7 |
| RNA-Seq data analysis | Circular RNA identification | Transcriptomics | 10.8 |
| Small RNA sequencing data analysis | microRNA quantification | Transcriptomics | 11.1 |
| Small RNA sequencing data analysis | microRNA prediction | Transcriptomics | 11.2 |
| CAGE-seq data analysis | TSS identification | Transcriptomics | 12.1 |
| 3' end-seq data analysis | PAS (polyadenylation site) identification | Transcriptomics | 13.1 |
| Nanopore RNA sequencing data analysis | Isoform expression | Transcriptomics | 14.1 |
| PacBio RNA sequencing data analysis | Isoform expression | Transcriptomics | 15.1 |
| CLIP-seq data analysis | Identify protein-RNA crosslink sites | Transcriptomics | 16.1 |
| RIP-seq data analysis | Find enriched genes bounded by RBP | Transcriptomics | 16.2 |
| Ribo-seq data analysis | Identify translated ORFs | Transcriptomics | 17.1 |
| single-cell RNA-seq data analysis | Cell clustering from fastq data | Transcriptomics | 18.1 |
| single-cell RNA-seq data analysis | Find differentially expressed genes based on count matrix | Transcriptomics | 18.2 |
| single-cell RNA-seq data analysis | Find marker genes based on count matrix | Transcriptomics | 18.3 |
| single-cell RNA-seq data analysis | Cell clustering and visualization | Transcriptomics | 18.4 |
| Spatial transcriptomics | Neighborhood enrichment analysis | Transcriptomics | 19.1 |
| Spatial transcriptomics | Single-cell mapping | Transcriptomics | 19.2 |
| Mass spectrometry data analysis | Protein expression quantification | Proteomics | 20.1 |
| Mass spectrometry data analysis | Metabolites quantification | Metabolomics | 21.1 |

Over the past few months, the rapid advancement of Large Language Models (LLMs) [57] has raised substantial expectations for the enhancement of scientific research, particularly in the field of biology [58], [59], [60]. These advancements hold promise for applications such as disease diagnosis [61], [62], [63], [64], drug discovery [65], and all. In the realm of bioinformatics, LLMs, such as ChatGPT, also demonstrate immense potential in tasks related to bioinformatics education [66] and code generation [67]. While researchers have found ChatGPT to be a valuable tool in facilitating bioinformatics research, such as data analysis, there remains a strong requirement for human intervention in the execution process. ChatGPT shows sensitivity to the nuances of user queries, resulting in diverse responses based on the prompts, which is the reason why prompt engineering is getting huge attention [68]. Given the specialized nature of bioinformatics tools, ChatGPT is also susceptible to potential issues, such as misinterpreting parameters, errors in software utilization, and other bugs that may arise during code generation. Users may encounter the necessity for ongoing engagement with ChatGPT, involving a continuous cycle of inquiry, code generation, execution, and debugging to ensure desired performance. AutoGPT [69], as a recently developed, advanced, and experimental open-source autonomous AI agent, has the capacity to string together LLM-generated “thoughts” to autonomously achieve user-defined objectives. Nevertheless, given the intricate and specialized nature of bioinformatics tasks, such as specialized software, the direct application of AutoGPT in this field still presents significant challenges. Notably, it faces difficulties in effectively managing the intricate software requirements of bioinformatics, encompassing tasks such as installation, software calls, and parameter settings.

In this study, we introduce Automated Bioinformatics Analysis (AutoBA), an autonomous AI agent tailored for comprehensive and conventional multi-omic analyses. AutoBA simplifies user interactions to just three inputs: data path, data description, and the final objective. This powerful tool autonomously proposes analysis plans, generates code, executes codes, and conducts subsequent data analysis by using our well-designed prompts. We implemented AutoBA as open-source software that offers multiple LLM backends, with options for both online and local usage, prioritizing data security and user privacy (Fig. 1). To show the reliability of AutoBA, we tested it in a large number of real-world multi-omic analysis scenarios (Fig. 2). AutoBA, serving as an AI agent tailored for bioinformatics data analysis, could address the surging demand for streamlined multi-omics data analysis, mitigate the financial challenges associated with bioinformatics training, and cater to diverse customization requirements. In summary, AutoBA is the first agent of this kind and represents a significant leap in the application of Large Language Models (LLMs) and automated AI agents within the domain of bioinformatics, highlighting their potential to accelerate future research in this field.

**Fig. 1.**
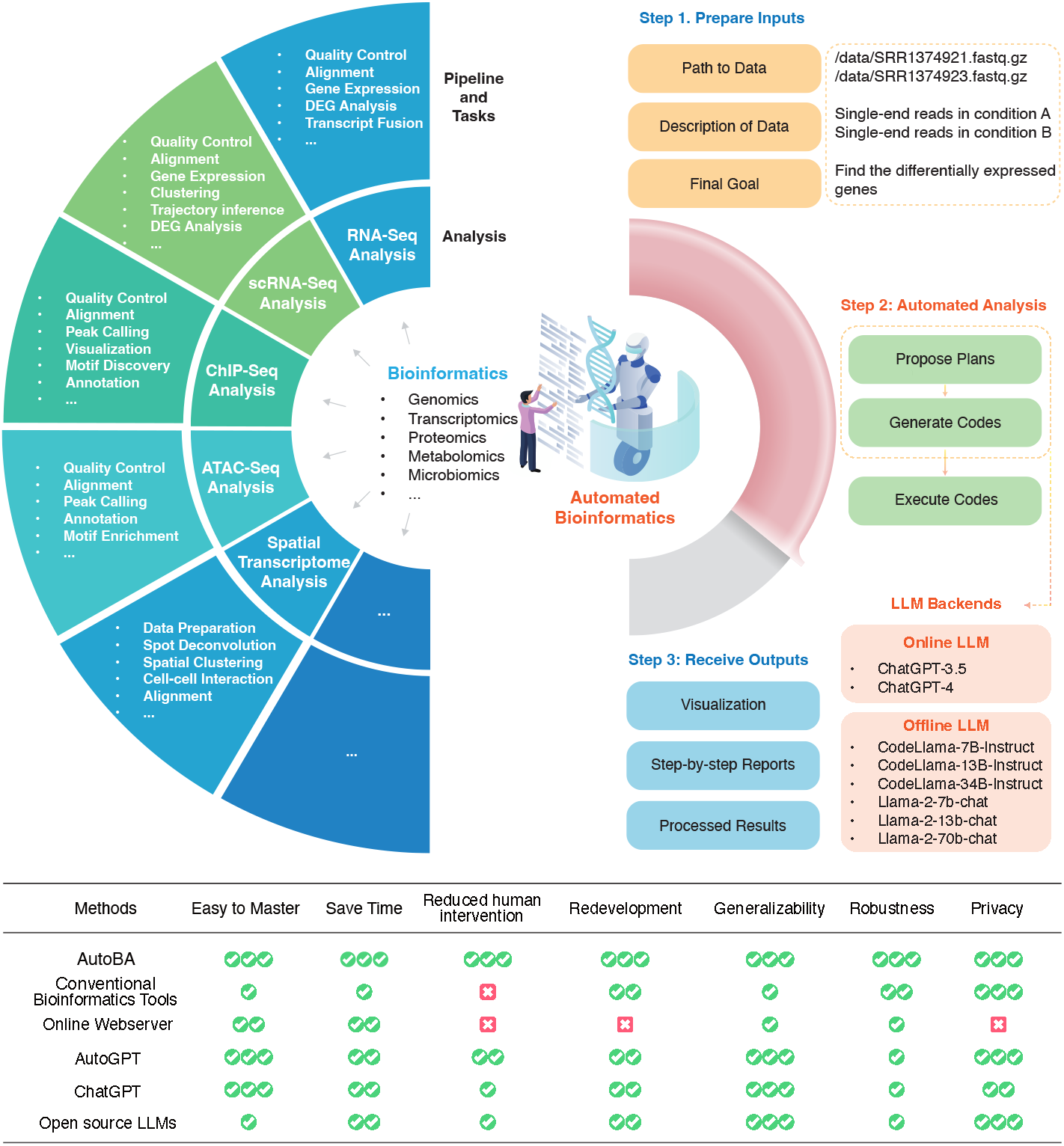
Design of AutoBA. AutoBA stands as the first autonomous AI agent meticulously crafted for conventional multi-omic analyses. Remarkably user-friendly, AutoBA simplifies the analytical process by requiring minimal user input, including data path, data description, and the final objective, while delivering detailed step-by-step plans for various bioinformatics tasks. With these inputs, it autonomously proposes analysis plans, generates code, executes codes, and conducts subsequent data analysis by using our well-designed prompts. AutoBA was implemented as open-source software that offers multiple LLM backends, with options for both online and local deployment, prioritizing data security and user privacy and offering a streamlined and efficient solution for bioinformatics tasks. Step 1 and Step 3 require human intervention, while Step 2 requires no human intervention. The table presents a qualitative comparison of AutoBA against other methods, assessing attributes such as user-friendliness, time efficiency, diminished human intervention (degree of automation), ease of redevelopment, generality, robustness, and privacy considerations.

**Fig. 2.**
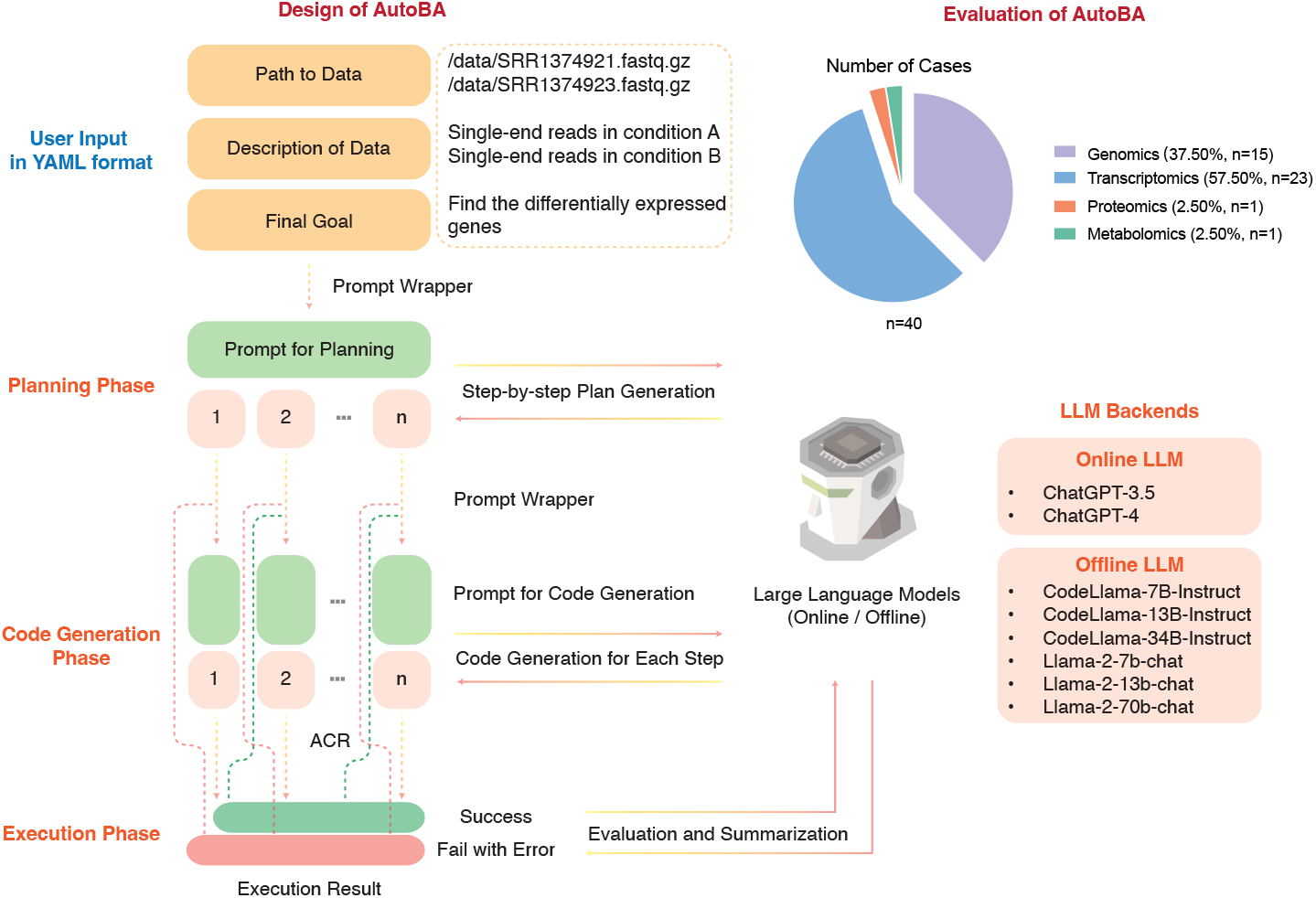
Detailed workflow design and evaluation of AutoBA. Pie chart indicates the number of all cases used for validating AutoBA.

## 2 Methods

### 2.1 The overall framework design of AutoBA

AutoBA is the first autonomous AI agent tailor-made for conventional multi-omic analyses. As illustrated in Fig. 1, conventional bioinformatics typically entails the use of pipelines to analyze diverse data types such as WGS, WES, RNA-seq, single-cell RNA-seq, ChIP-seq, ATAC-seq, spatial transcriptomics, and more, all requiring the utilization of various tools. Users are traditionally tasked with selecting the appropriate tools based on their specific analysis needs. In practice, this process involves configuring the environment, installing software, writing code, and debugging, which are time-consuming and labor-intensive.

With the advent of AutoBA, this labor-intensive process is revolutionized. Users are relieved from the burden of dealing with multiple software packages and need only provide three key inputs in YAML format: the data path (e.g., */data/SRR1374921*.*fasta*.*gz*), data description (e.g., *single-end reads in condition A*), and the ultimate analysis goal (e.g., *identify differentially expressed genes*). AutoBA takes over by autonomously analyzing the data, generating comprehensive step-by-step plans, composing code for each step, executing the generated code, and conducting in-depth analysis. Depending on the complexity and difficulty of the tasks, users can expect AutoBA to complete the tasks within a matter of minutes to a few hours, all without the need for additional human intervention (Table 1 and Fig. 2).

### 2.2 Prompt engineering of AutoBA

To initiate AutoBA, users provide three essential inputs: the data path, data description, and the previously mentioned analysis objective. AutoBA comprises three distinct phases: the planning phase, the code generation phase, and the execution phase as shown in Step 2 of Fig. 1. During the planning phase, AutoBA meticulously outlines a comprehensive step-by-step analysis plan. This plan includes details such as the software name and version to be used at each step, along with guided actions and specific sub-tasks for each stage. Subsequently, in the code generation phase, AutoBA systematically follows the plan and generates codes for sub-tasks, which entails procedures like configuring the environment, installing the necessary software, and writing code. Then, in the execution phase, AutoBA executes the generated code. In light of this workflow, AutoBA incorporates two distinct prompts: one tailored for the planning phase and the other for the code generation phase. Intensive experiments have shown that these two sets of prompts are essential for the proper functioning of AutoBA in automated bioinformatics analysis tasks.

The prompt for both the planning phase and the code generation phase are displayed in the supplementary information. In both prompt designs, the term *blacklist* pertains to the user’s personalized list of prohibited software. The current default blacklist contains several tools that frequently caused errors during our testing processes. Meanwhile, *data list* encompasses the inputs necessary for AutoBA, encompassing data paths and data descriptions. The term *current goal* serves as the final objective during the planning phase and as the sub-goal in the execution phase, while *history summary* encapsulates AutoBA’s memory of previous actions and information.

### 2.3 Memory management of AutoBA

A memory mechanism is incorporated within AutoBA to enable it to generate code more effectively by drawing from past actions, thus avoiding unnecessary repetition of certain steps. AutoBA meticulously logs the outcome of each step in a specific format, and all these historical records become part of the input for the subsequent prompt. In the planning phase, memories are structured as follows: “Firstly, you provided input in the format ‘file path: file description’ in a list: *<*data list*>*. You devised a detailed plan to accomplish your overarching objective. Your overarching goal is *<*global goal*>*. Your plan involves *<*tasks*>*.” In the code generation phase, memories follow this format: “Then, you successfully completed the task: *<*task*>* with the corresponding code: *<*code*>*.”

### 2.4 Automatic code repair of AutoBA

AutoBA incorporates an automatic code repair (ACR) module designed to streamline the debugging process and enhance the reliability of generated code. During the code execution phase, AutoBA identifies errors from the output stream called standard error (stderr). Once an error is detected, these detected errors will be integrated into the prompt for code regeneration, ensuring a repetitive cycle until the generated code successfully executes without errors.

### 2.5 Evaluation of AutoBA

The results produced by AutoBA undergo thorough validation by bioinformatics experts. This validation process encompasses a comprehensive review of the proposed plans, generated codes, execution of the code, and confirmation of the results for accuracy and reliability. AutoBA’s development and validation are built upon a specific environment and software stack, which includes Ubuntu version 18.04, Python 3.10.0, and openai version 0.27.6. These environment and software specifications form the robust foundation for AutoBA’s functionality in the field of bioinformatics, ensuring its reliability and effectiveness. To further assess the usability of AutoBA, we conducted a comparative analysis involving the following methods: 1) AutoBA (w/o ACR, online with ChatGPT-4), 2) AutoBA (with ACR, online with ChatGPT-4), 3) AutoBA (w/o ACR, offline with CodeLlama-34B-Instruct), 4) AutoBA (with ACR, offline with CodeLlama-34B-Instruct), 5) AutoGPT, 6) ChatGPT-3.5, 7) ChatGPT-4 and 8) CodeLlama-34B-Instruct. Given that prompt engineering and workflow design is a distinctive innovation of AutoBA, during the evaluation of AutoGPT, ChatGPT-3.5, ChatGPT-4, and CodeLlama-34B-Instruct, we emulated user behavior by utilizing a generalized and uniform prompt as shown in the supplementary information.

### 2.6 Online and local LLM backends of AutoBA

AutoBA offers several versions of LLM backends, including online backends based on ChatGPT-3.5 and ChatGPT-4, and local LLMs, including CodeLlama-7B-Instruct, CodeLlama-13B-Instruct, CodeLlama-34B-Instruct [70], Llama-2-7b-chat, Llama-2-13b-chat and Llama-2-70b-chat [71].

### 2.7 Security and safety of AutoBA

AutoBA incorporates a sandbox mode to establish a secure and isolated environment for conducting analyses. This mode encapsulates the analysis processes, effectively shielding the underlying system from potential threats. Meanwhile, AutoBA imposes restrictions on system commands throughout the execution phase, thereby reducing the risk of malicious commands being executed within the environment. Additionally, AutoBA leverages Docker containerization, introducing an extra layer of security to further fortify the overall system integrity. Furthermore, Docker containerization simplifies the installation process, contributing to a reduction in learning costs for users. A workstation with 252 GB RAM, 112 CPU cores, and 1 Nvidia A100 GPU was adopted for all experiments. AutoBA was developed based on Python3.10 and CUDA12.0. A detailed list of dependencies could be found in our code availability. The online version operates without the need for a GPU, while the offline version requires GPU support (7B: 12.55GB, 13B: 24GB, 34B: 63GB, 70B: 74GB).

## 3 Results

### 3.1 AutoBA proposes detailed analysis plans for tasks

AutoBA offers a robust capability to generate a highly detailed and customized analysis plan, leveraging the user’s input, which encompasses critical elements such as data paths, data descriptions, and objective descriptions.

As an example, in Fig. 3, the user supplied four RNA-Seq samples: two from the LoGlu group (SRR1374921.fastq.gz and SRR1374922.fastq.gz, mouse pancreatic islets cultured at low ambient glucose) and two from the HiGlu group (SRR1374923.fastq.gz and SRR1374924.fastq.gz, mouse pancreatic islets cultured at high ambient glucose) from Benner et al.’s paper [72]. Additionally, the user also provided the mouse reference genome (mm39.fa) and genome annotation (mm39.ncbiRefSeq.gtf). The primary objective of this case was to identify differentially expressed genes between the two data groups. Using textual inputs only, AutoBA generated a detailed, step-by-step analysis plan during the planning phase, as outlined below:

**Fig. 3.**
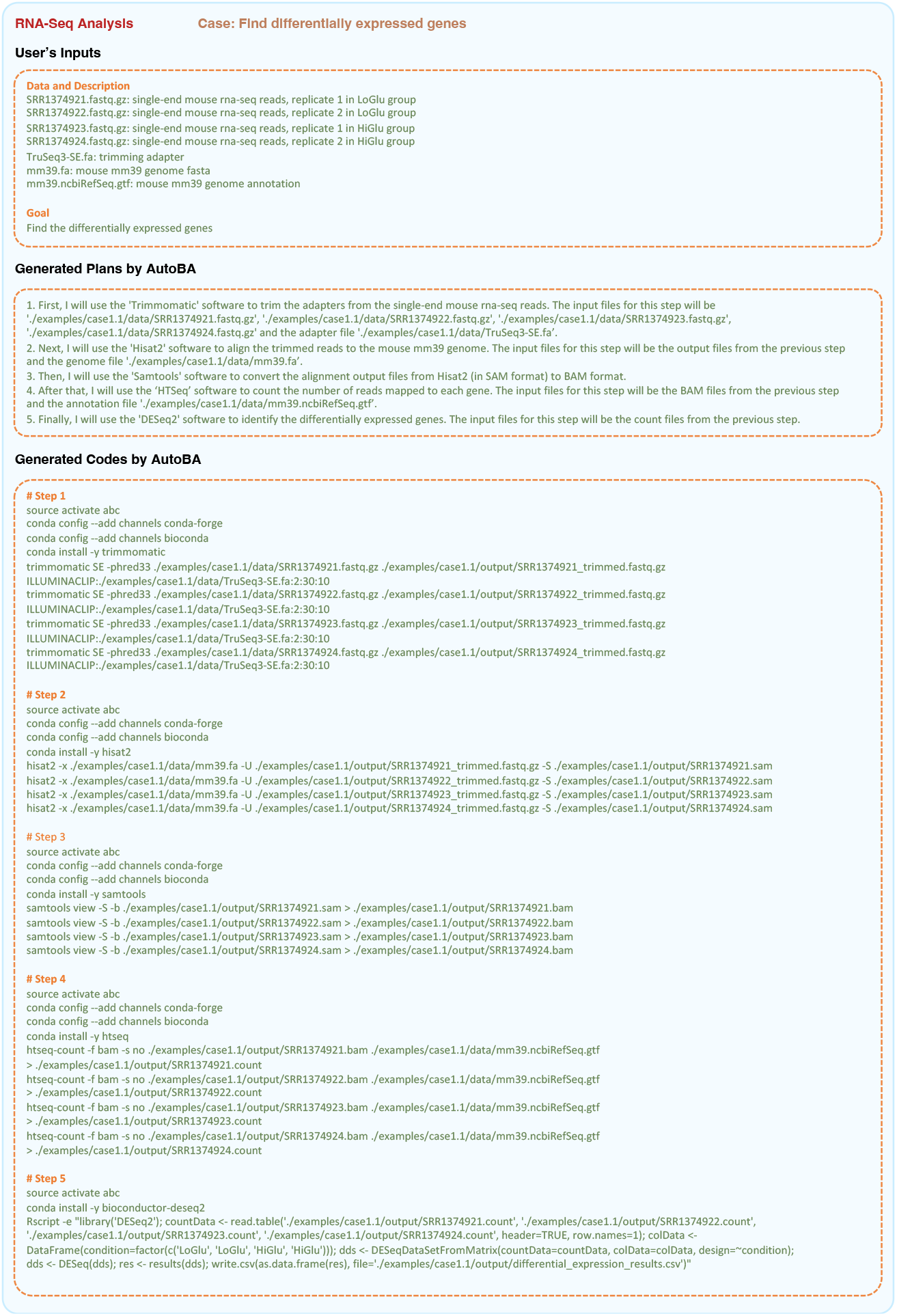
Example of applying AutoBA to find differentially expressed genes with RNA-Seq data. In this case, the user supplied four RNA-seq datasets, comprising two from the LoGlu group and two from the HiGlu group. The primary objective of this analysis was to identify differentially expressed genes across the two datasets.

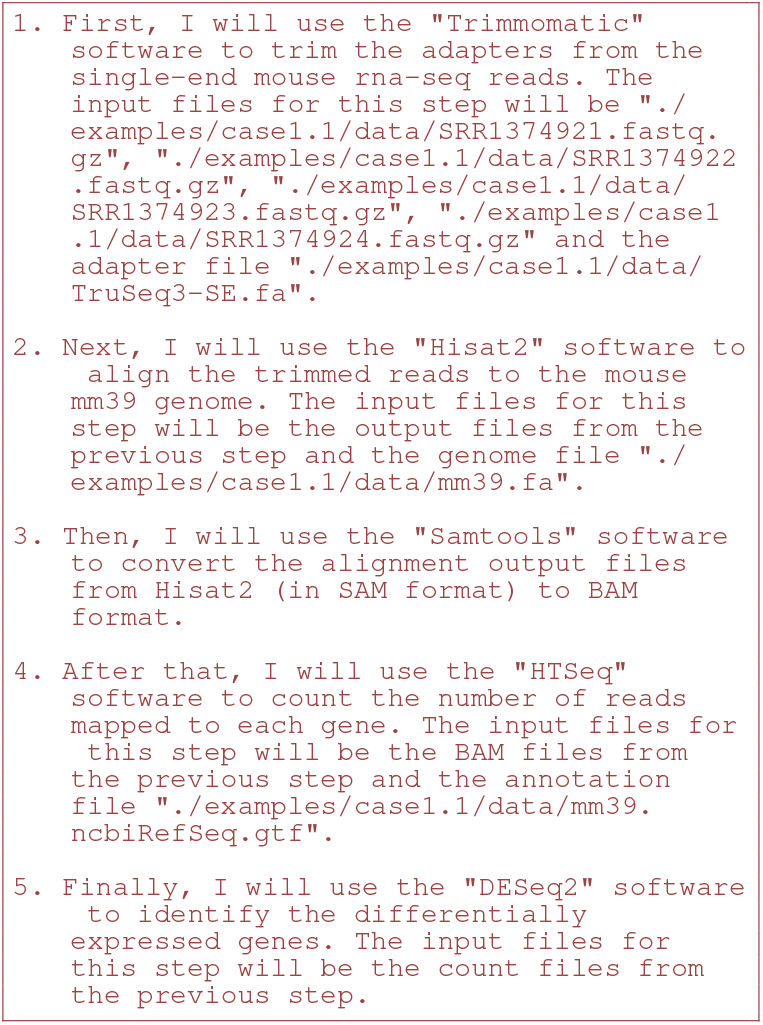

Within each step of the plan outlined above, AutoBA provides precise instructions regarding the required software, including names like Trimmomatic, Hisat2, Samtools, HTSeq, and DESeq2, along with clear sub-tasks for each analytical stage. This level of tailored planning ensures that the analysis process aligns precisely with the user’s objectives, promoting both efficiency and accuracy in data processing and results generation.

### 3.2 AutoBA generates precise codes for sub-tasks

During the code generation phase, AutoBA generates code in bash format for every sub-task of the plan established in the planning phase. These scripts encompass environment setup, software installation, and tailored code for software utilization. Parameters and data paths specific to the software are meticulously incorporated. As exemplified in Fig. 3, the preliminary phase of the differentially expressed genes (DEG) analysis constitutes the essential process of adapter trimming, an indispensable preprocessing step in the context of raw RNA-Seq data. Within this critical step, AutoBA automatically generated code, including activating the conda environment, installing software packages, and calling software to analyze data as shown below:

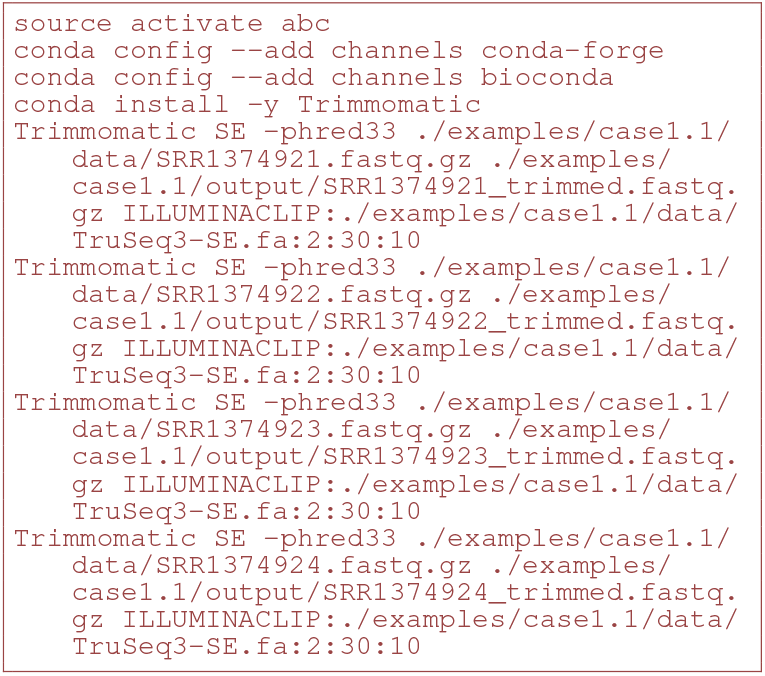

The generated code further underwent a meticulous and thorough validation process, which included a manual verification and execution performed by experienced and proficient bioinformaticians, as meticulously documented in Table 2 and Supplementary information. This critical validation step ensures the accuracy and reliability of the code, reaffirming the robustness of AutoBA.

**TABLE 2.** Summary of AutoBA (w/o ACR) generated results evaluated by bioinformatics experts. The table presents an assessment conducted by bioinformatics experts on the analysis plan proposed by AutoBA, along with the generated codes and the code execution. If the evaluation passes, it is displayed as success, while instances of failure are accompanied by detailed explanations of the specific reasons for the failure. Additionally, we provide a summary of the software tools automatically chosen by AutoBA for each case, as well as the total time taken to generate the corresponding code.

| Case ID | Propose Plans | Generate Codes | Execute Codes | Tools Used | Time Cost<br>(without Executing Codes)<br>in Minutes |
| --- | --- | --- | --- | --- | --- |
| 1.1 | Success | Success | Success | FastQC, Trimmomatic, SPAdes, QUAST | 3 |
| 2.1 | Success | Success | Success | FastQC, Trimmomatic, BWA, Samtools, GATK | 8 |
| 2.2 | Success | Success | Success | FastQC, Trimmomatic, BWA, Samtools, GATK, ensemble-vep | 8 |
| 2.3 | Success | Success | Success | FastQC, Trimmomatic, BWA, Samtools, GATK, manta | 18 |
| 2.4 | Success | Success | Failed: pindel<br>it requires configuration file | FastQC, Trimmomatic, BWA, Samtools, pindel, SnpEff | 6 |
| 3.1 | Success | Success | Success | FastQC, Trim Galore, Bowtie 2, Samtools, MACS2, BEDTools, IGV | 6 |
| 3.2 | Success | Success | Success | FastQC, Trim Galore, Bowtie2, MACS2, HOMER, MEME | 4 |
| 3.3 | Failed:<br>DESeq2 is not suitable for<br>peaks identified by MACS2 | - | - | FastQC, BWA, MACS, BEDTools, DESeq2, g:Profiler, R | 6 |
| 4.1 | Success | Success | Success | Trim Galore, Bismark, IGV | 9 |
| 5.1 | Success | Success | Failed:<br>(wrongly used BEDTools) | Trim Galore, BWA, Samtools, MACS2, BEDTools | 8 |
| 6.1 | Success | Success | Success | FastQC, Cutadapt, BWA, MACS2, IGV, GREAT | 5 |
| 7.1 | Success | Success | Success | FastQC, BEDTools, Samtools, Bowtie 2, R | 6 |
| 8.1 | Success | Success | Failed: racon<br>medaka<br>wrongly used the parameters | canu, Minimap2, Racon, Flye, Medaka, Bandage | 7 |
| 8.2 | Failed:<br>cannot find a correct pipeline | - | - | Minimap2, Samtools, trf | 7 |
| 9.1 | Success | Failed: install the wrong tool,<br>pb-falcon rather than falcon | - | Canu, FALCON, Quiver, MUMmer, | 7 |
| 10.1 | Success | Success | Success | FASTQC, Trimmomatic, HISAT2, htseq, DESeq2 | 5 |
| 10.2 | Success | Success | Success | FASTQC, Trimmomatic, HISAT2, htseq, DESeq2, gprofileR | 5 |
| 10.3 | Success | Success | Success | gunzip, HISAT2, fusioncatcher, gffcompare | 6 |
| 10.4 | Success | Success | Success | Trim Galore, HISAT2, Samtools, StringTie | 5 |
| 10.5 | Success | Success | Success | Trimmomatic, HISAT2, Samtools, StringTie, featureCounts, rMATs | 6 |
| 10.6 | Success | Failed: DaPars<br>(not available in conda) | - | Trim Galore, HISAT2, StringTie, DaPars | 7 |
| 10.7 | Failed:<br>cannot find a correct pipeline | - | - | FastQC, Trimmomatic, HISAT2, Samtools, StringTie, ballgown, GATK | 7 |
| 10.8 | Success | Failed: CIRI2<br>(not available in conda) | - | Trim Galore, HISAT2, CIRI2, CIRIQuant | 5 |
| 11.1 | Success | Success | Success | Fastqc, Cutadapt, Bowtie, Samtools, subread/featureCounts, DESeq2, edgeR | 11 |
| 11.2 | Success | Success | Failed: conda of miRDeep2<br>is problematic | Fastqc, Cutadapt, Bowtie, Samtools, featureCounts, miRDeep2, DESeq2, edgeR | 11 |
| 12.1 | Success | Success | Success | Fastqc, Trimmomatic, HISAT2, HTSeq/htseq-count, CAGEr | 6 |
| 13.1 | Failed:<br>cannot find a correct pipeline | - | - | Trim Galore, HISAT2, StringTie, DaPars | 5 |
| 14.1 | Success | Success | Failed: prepDE.py no need<br>to run with 'python prepDE.py' | Minimap2, Samtools, StringTie, DESeq2 | 9 |
| 15.1 | Success | Success | Success | Minimap2, Samtools, StringTie, cufflinks | 5 |
| 16.1 | Success | Success | Failed: conda of Piranha is problematic | FastQC, Cutadapt, Bowtie2, Samtools, BEDTools, Piranha | 6 |
| 16.2 | Success | Success | Success | FastQC, Trim Galore, HISAT2, htseq, DESeq2 | 4 |
| 17.1 | Success | Success | Failed: not regular conda of ribotaper | FastQC, Trim Galore, HISAT2, Samtools, StringTie, RiboTaper | 7 |
| 18.1 | Success | Success | Success | Cell Ranger, Seurat | 5 |
| 18.2 | Success | Success | Success | Scanpy | 8 |
| 18.3 | Success | Success | Success | Scanpy | 6 |
| 18.4 | Success | Success | Success | Scanpy | 5 |
| 19.1 | Success | Success | Success | Squidpy, AnnData | 5 |
| 19.2 | Success | Success | Success | AnnData, Scanpy, Tangram | 3 |
| 20.1 | Success | Success | Success | proteowizard, OpenMS | 15 |
| 21.1 | Success | Success | Success | pymzml, pandas, numpy, scipy | 13 |
| #Success | 36 | 33 | 26 | - | - |

### 3.3 AutoBA adeptly manages similar tasks with robustness

In practical bioinformatics applications, even when researchers are working with similar data types, such as RNA-Seq, it’s noteworthy that analyses often manifest variations stemming from diverse sources. These variations are primarily attributed to disparities in the characteristics of input data and the distinct objectives pursued in the analytical process.

As exemplified in Case 10.1 (find differentially expressed genes), Case 10.2 (identify the top five down-regulated genes in HiGlu group), and Case 10.3 (predict fusion genes), when performing RNA-Seq analysis, users may have distinct final goals, necessitating adjustments in software and parameter selection during the actual execution. In comparison to case 10.1, AutoBA introduces an additional step in case 10.2, tailored for screening the top five differentially expressed genes to fulfill the user’s specific requirements as shown in the code below:

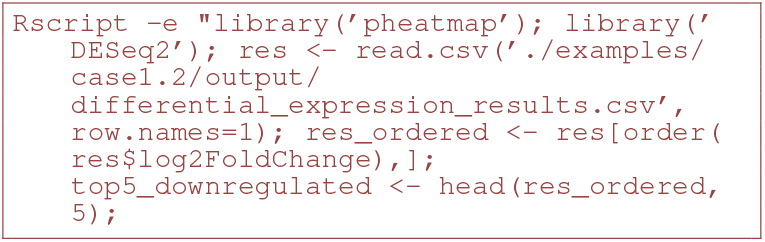

### 3.4 AutoBA adjusts analysis based on task and input data variations

Alignment is an essential step for bioinformatic analysis, for which multiple tools have been developed for distinct tasks. For instance, tools including STAR [73] and HISAT2 [74] designed for RNA-seq data analysis are splicing aware, which is efficient in identifying junction reads that map to two distal positions in the reference genome. Besides, long-read sequencing data from Pacific Bioscience (PacBio) and Oxford Nanopore Technology (ONT) also require specialized tools for the alignment, for which Minimap2 [75] is the most widely used method. Moreover, each read from single-cell sequencing data contains barcodes for UMI and cell labels, which needs to be integrated with the alignment. CellRanger is a popular software with this capacity. Therefore, bioinformatic analysis should use appropriate tools for the alignment based on the types of tasks. Interestingly, we found that AutoBA has learned this knowledge and can correctly employ the tool for the alignment (Fig. 4a).

For many bioinformatic analysis, multiple tools are available but require different conditions of inputs. For instance, to identify structural variations from tumor WGS/WES data, the method “manta” [76] can handle the analysis against the matched normal. On the other hand, tools like “Pindel” [77] that relies on the detection of breakpoints with the reference genome, only conduct analysis on the tumor samples. We found that AutoBA can automatically select “manta” when the matched normal samples were provided and correctly utilized the parameters “–normalBam” and “–tumorBam”. However, if only the tumor samples were provided in the input data, AutoBA will select “Pindel” for the analysis (Fig. 4b). These results suggest that AutoBA learned the requirements of different bioinformatic tools and is capable of selecting appropriate tools based on different conditions of the input data.

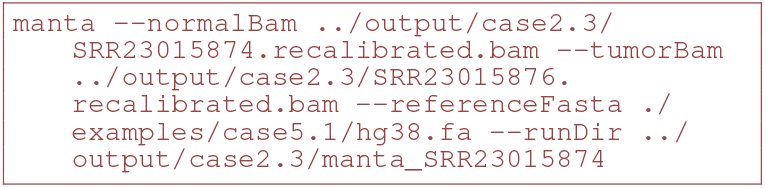

### 3.5 Apply AutoBA to a variety of conventional multiomic analyses scenarios

To evaluate the robustness of AutoBA, we conducted assessments involving a total of 40 cases spanning four distinct types of omics data: genomics, transcriptomics, proteomics, and metabolomics as shown in Table 1 and Supplementary information.

All cases underwent an independent analysis process conducted by AutoBA and were subsequently subjected to validation by experienced bioinformatics experts. The collective results underscore the versatility and robustness of AutoBA across a spectrum of multi-omics analysis procedures in the field of bioinformatics as shown in Table 2. AutoBA demonstrates its capability to autonomously devise novel analysis processes based on varying input data, showcasing its adaptability to diverse input data and analysis objectives with a success rate of 90% (36 out of 40) for proposing plans, 82.5% (33 out of 40) for generating codes to obtain and install appropriate tools, and 65% (26 out of 40) for automated end-to-end analysis. With the incorporation of the ACR module, AutoBA demonstrates enhanced robustness, with the same success rate of 90% (36 out of 40) for proposing plans, but a higher success rate of 87.5% (35 out of 40) for generating codes to obtain and install appropriate tools, and 87.5% (35 out of 40) for automated end-to-end analysis. Compared to the online version, the local version showed a slight decline in performance as shown in Fig. 4d.

**Fig. 4.**
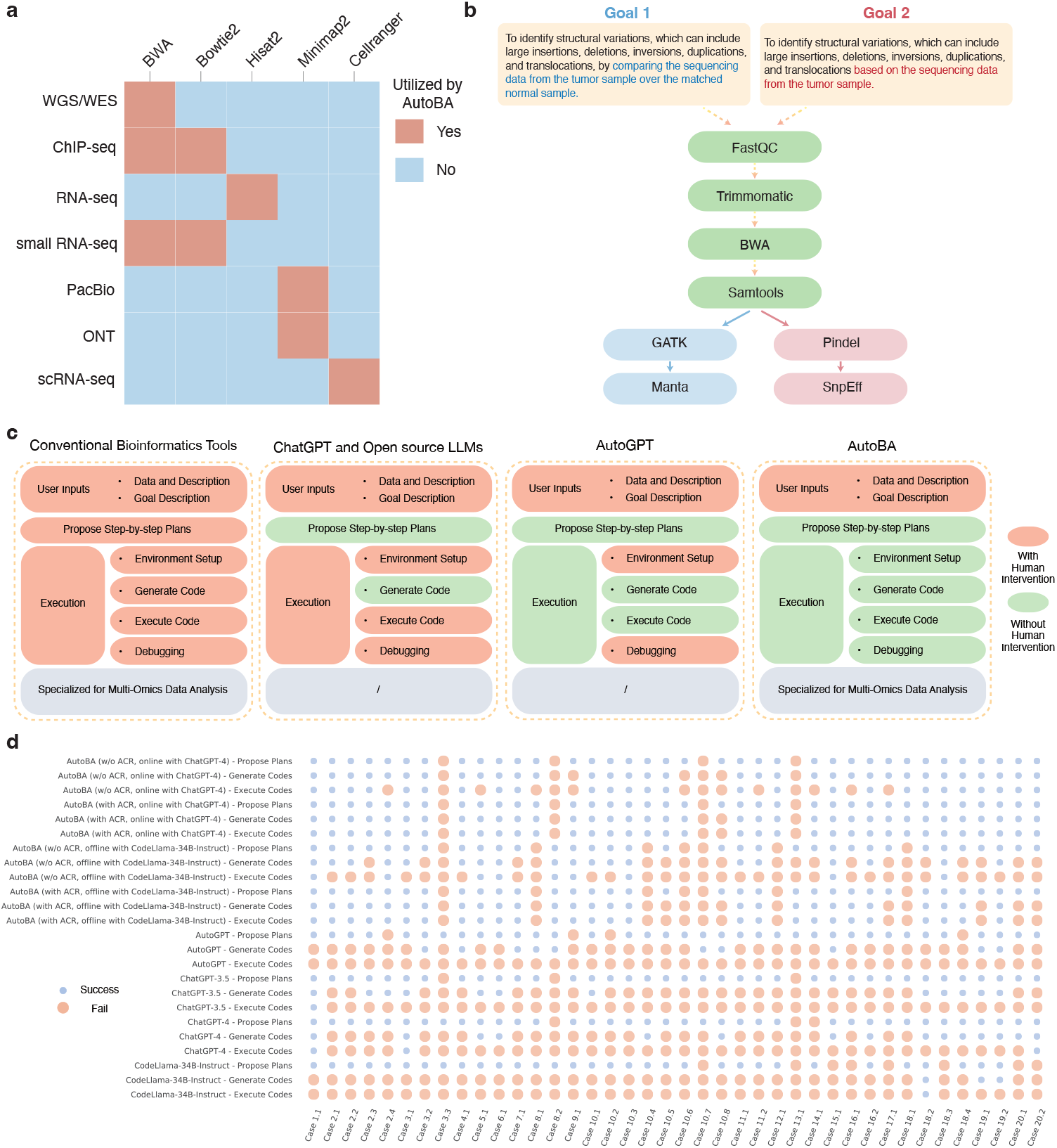
Results of AutoBA and the comparison with other methods. **a** Heatmap illustrating options of utilizing different alignment tools for multiple tasks planned by AutoBA. **b** AutoBA utilizes the tools for identifying structure variations in tumor samples with or without the matched normal samples. The highlight shows the difference between Goal 1 and Goal 2. **c** Conceptual comparison of AutoBA with other methods in terms of human intervention. Orange indicates the need for human intervention, while green signifies an absence of human intervention (fully automated process). **d** Evaluation of results generated by various methods by manually checking and executing codes and comparing them to standard analysis pipelines. Orange indicates a failure, and blue indicates a success.

### 3.6 AutoBA reduces human intervention and increases robustness compared to other methods

As shown in Fig. 4

To show the robustness of AutoBA, we further conducted a comprehensive comparison of eight methods, including 1) AutoBA (w/o ACR, online with ChatGPT-4), 2) AutoBA (with ACR, online with ChatGPT-4), 3) AutoBA (w/o ACR, offline with CodeLlama-34B-Instruct), 4) AutoBA (with ACR, offline with CodeLlama-34B-Instruct), 5) AutoGPT, 6) ChatGPT-3.5, 7) ChatGPT-4 and 8) CodeLlama-34B-Instruct, across all 40 cases, as illustrated in Fig. 4d. AutoBA showed better performance in comparison to AutoGPT (90% for proposing plans, 25% for generating codes to obtain and install appropriate tools, and 0% for automated end-to-end analysis), ChatGPT-3.5 (92.5% for proposing plans, 30% for generating codes to obtain and install appropriate tools, and 2.5% for automated end-to-end analysis), ChatGPT-4 (92.5% for proposing plans, 37.5% for generating codes to obtain and install appropriate tools, and 7.5% for automated end-to-end analysis), and CodeLlama-34B-Instruct (80% for proposing plans, 7.5% for generating codes to obtain and install appropriate tools, and 2.5% for automated end-to-end analysis).

## 4 Discussion

To our knowledge, AutoBA is the first autonomous AI agent tailored explicitly for conventional multi-omic analyses for omics data. AutoBA streamlines the analytical process, requiring minimal user input while providing detailed step-by-step plans for various bioinformatics tasks (Video S1). The results of our investigation reveal that AutoBA excels in accurately handling a diverse array of omics analysis tasks, such as RNA-seq, scRNA-seq, ChIP-seq, spatial transcriptomics, and so on. One of the key strengths of AutoBA is its adaptability to variations in analysis objectives. As demonstrated in the cases presented, even with similar data types, such as RNA-Seq, users often have distinct goals, necessitating modifications in software and parameter selection during execution. AutoBA effectively accommodates these variations, allowing users to tailor their analyses to specific research needs without compromising accuracy. Furthermore, AutoBA’s versatility is highlighted by its ability to self-design new analysis processes based on differing input data. This autonomous adaptability makes AutoBA a valuable tool for bioinformaticians working on novel or unconventional research questions, as it can adjust its approach to the unique characteristics of the data.

Online bioinformatics analysis platforms are currently in vogue, but they often necessitate the uploading of either raw data or pre-processed statistics by users, which could potentially give rise to privacy concerns and data leakage risks. In contrast, AutoBA addresses these privacy issues by offering both online version and local version. When utilizing the online version of AutoBA with ChatGPT, data uploads are unnecessary, requiring only descriptive information in natural language as specified in our prompt design. This information is limited in terms of private details. In comparison, the local version of AutoBA provides the highest level of privacy protection, as it operates on local backends and eliminates the need to share any information with third parties. Moreover, AutoBA showcases its adaptability in sync with emerging bioinformatics tools, with LLM seamlessly incorporating these latest tools into the database. Furthermore, AutoBA is inclined towards selecting the most popular analytical frameworks or widely applicable tools in the planning phase, underscoring its robustness. Another distinguishing feature is AutoBA’s transparent and interpretable execution process. This transparency allows professional bioinformaticians to easily modify and customize AutoBA’s outputs, leveraging AutoBA to expedite the data analysis process.

Given that classical bioinformatic analysis encompasses a far broader spectrum of tasks and challenges than the 40 cases studied in this work (Table 1 and 2), it is essential to conduct more real-world applications by our potential users to further comprehensively validate the robustness of AutoBA. We found that a large proportion (36%, 5 out of 14) of failed cases in executing code is due to the tools in conda being problematic, not in a regular form (end with .sh, .pl et al), or requiring an edited config file, suggesting a demand for more standard bioinformatics tools. Furthermore, taking into account the timeliness of the training data used for large language models, it’s important to note that some of the most recently proposed methods in bioinformatics may still pose challenges in automatically generating code by AutoBA. Therefore, a future endeavor to train an up-to-date large language model explicitly tailored for bioinformatics can significantly enhance AutoBA’s ability to maintain up-to-date code generation capabilities. Nevertheless, AutoBA represents a significant advancement in the field of bioinformatics, offering a user-friendly, efficient, and adaptable solution for a wide range of omics analysis tasks. Its capacity to handle diverse data types and analysis goals, coupled with its robustness and adaptability, positions AutoBA as a valuable asset in the pursuit of accelerating bioinformatics research. We anticipate that AutoBA will find extensive utility in the scientific community, supporting researchers in their quest to extract meaningful insights from complex biological data.

## 5 Data availability

The RNA-seq dataset could be downloaded from Sequence Read Archive (SRA) with IDs: SRR1374921, SRR1374922, SRR1374923, and SRR1374924. The dataset for case 1.3 could be downloaded from https://github.com/STAR-Fusion/STAR-Fusion-Tutorial/wiki. The scRNA-seq dataset could be downloaded from http://cf.10xgenomics.com/samples/cell-exp/1.1.0/pbmc3k/pbmc3k filtered gene bc matrices.tar.gz. The ChIP-seq dataset could be downloaded with IDs: SRR620204, SRR620205, SRR620206, and SRR620208. The Spatial Transcriptomics dataset could be downloaded from https://doi.org/10.5281/zenodo.6334774. The CAGE-seq dataset could be downloaded from SRA with IDs: SRR11351697, SRR11351698, SRR11351700, and SRR11351701. The 3’end-seq dataset could be downloaded from SRA with IDs: SRR17422754, SRR17422755, SRR17422756, and SRR17422757. The CLIP-seq dataset could be downloaded from ENCODE (https://www.encodeproject.org) with IDs: ENCLB742AYH and ENCLB770EDJ. The Ribo-seq data could be downloaded from SRA with IDs: RR12354645 and RR12354646. The raw single-cell RNA sequencing data could be downloaded from 10X genomics. The PacBio long-read sequencing data could be downloaded from SRA with IDs: SRR19552218 and SRR19785215. The small RNA-seq data could be downloaded from the previous study [78].

## 5 Code availability

The AutoBA software is publicly available at https://github.com/JoshuaChou2018/AutoBA. The Docker version of AutoBA is available at https://hub.docker.com/r/joshuachou666/autoba

## 6 Credit author statement

Conceptualization: J.Z. and X.G. Design: J.Z., B.Z. and X.G. Code implementation: J.Z. Application: J.Z., B.Z., X.C., H.L., C.X., W.H. Drafting of the manuscript: J.Z. and B.Z. Critical revision of the manuscript for important intellectual content: J.Z., B.Z., X.X., S.C., X.G. Supervision: J.Z. and X.G. Funding acquisition: X.G.

## 8 Acknowledgements

Juexiao Zhou, Bin Zhang, Xiuying Chen, Haoyang Li, Xiaopeng Xu, Siyuan Chen, Wenjia He, Chencheng Xu and Xin Gao were supported in part by grants from the Office of Research Administration (ORA) at King Abdullah University of Science and Technology (KAUST) under award number FCC/1/1976-44-01, FCC/1/1976-45-01, REI/1/5202-01-01, REI/1/5234-01-01, REI/1/4940-01-01, RGC/3/4816-01-01, REI/1/0018-01-01, REI/1/5414-01-01, REI/1/5289-01-01, and REI/1/5404-01-01

